# Darwinian selection of host and bacteria supports emergence of Lamarckian-like adaptation of the system as a whole

**DOI:** 10.1101/093120

**Authors:** Dino Osmanovic, David A Kessler, Yitzhak Rabin, Yoav Soen

## Abstract

**Background:** The relatively fast selection of symbiotic bacteria within hosts and the potential transmission of these bacteria across generations of hosts raise the question of whether interactions between host and bacteria support emergent adaptive capabilities beyond those of germ-free hosts.

**Results:** To investigate possibilities for emergent adaptations that may distinguish composite host-microbiome systems from germ-free hosts, we introduce a population genetics model of a host-microbiome system with vertical transmission of bacteria. The host and its bacteria are jointly exposed to a toxic agent, creating a toxic stress that can be alleviated by selection of resistant individuals and by secretion of a detoxification agent (“detox”). We show that toxic exposure in one generation of hosts leads to selection of resistant bacteria, which in turn, increases the toxic tolerance of the host’s offspring. Prolonged exposure to toxin over many host generations promotes additional form of emergent adaptation due to selection of hosts based on detox capabilities of their bacterial community as a whole (as opposed to properties of individual bacteria).

**Conclusions:** These findings show that interactions between pure Darwinian selections of host and its bacteria can give rise to emergent adaptive capabilities, including Lamarckian-like adaptation of the host-microbiome system.

## Background

Evolutionary adaptations are commonly thought to be driven by genetic mutations occurring on a timescale of many generations. Selection of individuals with rare beneficial mutations and transmission of the mutations across generations can then support adaptive evolution of the population. The exclusive focus on mutations that rarely occur during the lifetime of an individual has recently been expanded [1-7] to mechanisms supporting various forms of non-Mendelian inheritance, including: transgenerational epigenetic phenomena [8-11], genome editing and mobility systems [12, 13], niche construction [14] and transmission of symbiotic microorganisms [3, 5, 6, 15-19]. The case of symbiotic organisms may be of particular interest because of its broad relevance to animals and plants and the potential of host-microbe interactions to support adaptations that were traditionally considered impossible for hosts and bacteria on their own [3, 5, 17, 20, 21]. This is primarily due to a fundamental distinction between a composite, host-microbiome system and germ-free hosts, namely that the former undergoes intertwined selections, operating on different timescales: rapid selections of symbiotic microorganisms within the host and slower selection of that host (with its bacterial population). While the selection of each bacterium is governed by its individual traits, selection of the host depends jointly on the traits of the host and the properties of its bacterial community [20, 22-27]. This community can vary during the lifetime of the host and resident bacteria can be transferred across generations and/or between neighboring hosts [28-34]. Whether symbiosis between a host and microorganisms (collectively referred to as a holobiont [17] [35]) warrants a significant change to evolutionary thinking is currently under debate [36-39]. In particular, it is not clear whether the association between host and bacteria is tight enough to consider the holobiont as a unit of selection in evolution [36, 37] and whether transmission of bacteria across generations of hosts is stable enough to support non-traditional adaptive capabilities. To investigate the feasibility of emergent adaptations, we introduce a modeling framework that avoids speculative assumptions and relies instead on interactions between well-accepted Darwinian selection of host and resident bacteria. This allows us to study how particular types of interactions influence the adaptation of host and bacteria on a wide range of timescales. Our modelling approach builds on the traditional framework of population genetics [40, 41], but extends it in order to account for important considerations of host-microbiome systems that are not relevant for a population of germ-free hosts. In this model, we evaluate the adaptation of host and vertically-transmitted bacteria that are jointly exposed to a toxic agent. The exposure promotes Darwinian selections that occur on different timescales for host and bacteria. We find that the combined effect of these selections has profound implications. Among these, we show that the interaction between the selections of host and bacteria can give rise to an emergent, Lamarckian-like adaptation of the host-microbiome system within a single host generation. This effect is mediated by distinct modes of stress alleviation during a single host generation and it has a non-trivial dependence on the environmental conditions and on the characteristics of the system. Persistence of the exposure over timescales longer than a host generation promotes additional selection of hosts with bacterial communities that secrete higher average detox per bacterium (in contrast to selection of better fit individual bacteria, which takes place on much shorter timescales). This gives rise to a second mode of emergent adaptation that is independent of the Lamarckian-like adaptation within a single generation. In both cases, however, most of the adaptive benefit to the host is not attributable to changes in its own traits, but rather to alterations in the bacterial community. These alterations promote an increase in toxin tolerance which persists over periods longer than a host generation but shorter than typical evolutionary timescales.

## Results

### General considerations of the model

We consider the simplest case of a host-microbiome system in which every host is associated with a single species of symbiotic bacteria that is transmitted to the offspring with perfect fidelity. We take the generation time of a host to be much larger than for bacteria and we probabilistically determine the survival of each host and bacteria, according to the state of the organism at the end of their respective generation time (as detailed below). Each surviving host and bacterium gives rise to one offspring that inherits the traits of its parent, subject to a small random modification depending on a constant mutation rate, ***µ*** (no epigenetics is considered). The host and its bacterial community are jointly exposed to a toxin of concentration ***T***, thus creating a stress that impacts their survival probability of the host and each of its bacteria. This stress depends both on their intrinsic traits and on how they interact with one another. To investigate whether and how the coupling between the survival of host and bacteria could support non-traditional modes of adaptation, we consider broadly applicable types of interactions between host and bacteria. The mathematical representations of these interactions was chosen to simplify the identification and analysis of general effects which apply to many host-microbiome systems (as opposed to a model designed to fit a specific system).

We start by defining the toxic stress experienced by individual host and bacterium. Since this stress depends on the level of toxin, ***T***, and on the individual’s sensitivity to the toxin, ***x***, we define the instantaneous toxic stress for host and bacteria as ***S*_*H*_ = *x*_*H*_ *T*** and ***S*_*B*_ = *x*_*B*_ *T***, respectively. Accordingly, this stress can be alleviated by cell-intrinsic reduction in sensitivity and/or by secretion of a detoxifying agent, “detox” that reduces the toxic challenge (with or without associated cost).

Unlike in a germ-free system of hosts, a host in a composite host-microbiome system is influenced (and/or is dependent) on bacterial-derived nutrients and various other factors [42-44]. Exposure to toxin may therefore lead to physiological stress to the host due to a significant loss of bacteria. An indirect stress to the host can also be induced by factors that promote a significant excess of bacteria. We model these effects by introducing a physiological stress, ***S*_*ph*_**, which depends on deviations from a preferred size of the bacterial population. From the bacterial perspective, on the other hand, the host provides a niche (carrying capacity for bacteria). In the simpler case of free-living bacteria, the carrying capacity is typically modelled by a constant parameter, representing the extractable resources from an unchanging environment. The fixed niche assumption does not necessarily hold when the bacteria are accommodated inside a host which can modulate the size of the niche under stress [22]. Since we do not know in advance whether and how a host’s stress influences the number of bacteria that can be accommodated, we constructed a population model in which this influence is determined by natural selection.

Altogether, the model considers host-microbiome interactions that are mediated by: (i) mutual alleviation of toxic challenge via secretion of a detoxification agent (“detox”), (ii) dependence of the hosts’ well-being on the size of the bacterial population and (iii) modulation of the bacterial niche size by the stress state of the host.

### Model formulation

For each host and bacterium, we assign a probability of survival to reproduction, *P*_*H*_ and *P*_*B*_ respectively, defined as follows:

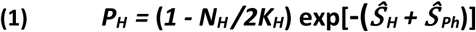

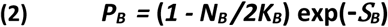

Here, ***N*_*H*_** and ***N*_*B*_** are the population sizes of hosts and bacteria per host, respectively, ***K*_*H*_** is the maximal number of hosts that can be supported by the external environment (carrying capacity for hosts) and ***K*_*B*_** is the number of bacteria that can be accommodated in the host (carrying capacity for bacteria). The toxic and physiological stress to the host, ***Ŝ*_*H*_ = <*S*_*H*_*>*_*t*_** and ***Ŝ*_*Ph*_ = *In*(<*N*_*B*_*>*_*t*_*/K*_*B*_^*0*^)** + (1 **- <*N*_*B*_*>*_*t*_*/K*_*B*_^*0*^)**, are defined respectively in terms of time averages of ***S*_*H*_** and ***N*_*B*_** over a host generation time (interval between host reproduction events; recall that the probability of survival is calculated only at the end of each generation). The physiological stress vanishes when the time-averaged bacterial population, **<*N*_*B*_ *>* _*t*_**, reaches a preferred size determined by a fixed parameter, ***K*_*B*_^*0*^**. The latter also sets an inverse scale (***1/K*_*B*_^*0*^**) for the negative impact of losing too many bacteria or having to support excess numbers of bacteria [43].

To test if selection might favor hosts that react to toxic stress by modulating the niche available for bacteria [45-48], we consider a population of hosts, each with a distinct dependence of the carrying capacity on the toxic stress of the host. For that, we define ***K*_*B*_** as:

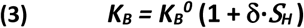

where **δ** is an evolvable trait, determining how the bacterial niche in the host is affected by the toxic stress it experiences. Since bacteria can affect this stress by secreting detox on a timescale shorter than a host generation, ***K*_*B*_** is jointly influenced by the host and the bacteria. To enable unbiased analysis of how ***K*_*B*_** changes in response to selection under exposure to toxin, we considered a starting population of hosts with a broad distribution of **δ**’s, symmetric around zero.

We assume that all the hosts and their bacteria are exposed, at time ***t***, to the same influx of active toxin, ***ϑ(t)***, applied instantaneously (i.e., in one bacterial generation, ***Δt***). This toxin can be neutralized by release of detox from the host and each of its bacteria [43, 48-50]:

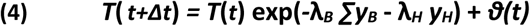

where ***y*_*H*_** and ***y*_*B*_** are the instantaneous amounts of detox secreted inside the host (by resident bacteria and the host itself) and **λ_*H*_** and **λ_*B*_** are the respective detoxification capacities of host and bacteria. We assume that all the bacteria of a given host benefit equally from the total amount of detox, regardless of their individual contributions to this total amount. The effect of having a cost associated with the secretion of detox by the bacteria is investigated in an extended version of the model (Supplementary Information).

The evolvable traits of the model (***x***, ***y*** and **δ**) are initially drawn from trait-specific distributions and are modified by the joint actions of mutation and selection. Surviving bacteria divide at every time step of the simulation (***Δt***), while the surviving hosts reproduce every **τ** generations of bacteria (so that the host generation time is **τ *Δt***). We consider the simplest reproduction model in which each of the surviving hosts and bacteria gives rise to one offspring that inherits the traits of its parent, subject to a small random modification depending on a constant mutation rate, ***µ***:

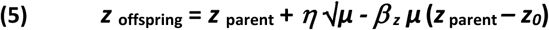

Here ***z*** corresponds to any of the evolving traits ***x***, ***y*** and **δ**, ***η*** is a standard Gaussian deviate with zero mean, and the parameters, ***z*_*0*_** and ***β* _*z*_** are trait-specific coefficients controlling the peak and width of the steady state distributions (specified in Methods). Note that 1/***β* _*z*_** sets a characteristic time for the distribution of a trait **z** to return to steady state, following an initial perturbation. The values of ***β* _*y*_** and ***β* _*δ*_** were chosen to support broad distributions of ***y*** and **δ**, respectively. To prevent a trivial solution in which all the individuals are completely insensitive to toxin, the sensitivity distribution (i.e. for ***z*** *=* ***x*_*H*_** and ***x*_*B*_**) is truncated at ***x*** = 0. We also avoid negative values of detox secretion by setting negative ***y*** values in Eq. 5 to zero. The remaining dynamic variables are updated in every generation of bacteria (***N*_*B*_**, ***T***, ***S*_*H*_**, ***S*_*B*_** and ***K*_*B*_**) and host (***N*_*H*_**). This study was based on an initial population of 32000 hosts (***N*_*H*_ *= K_H_*** = 32000) with 100 bacteria per host (***N*_*B*_ *= K*_*B*_^*0*^** = 100). The host generation time was set to **τ =** 100 bacterial generations and all the mutation rates were ***µ*** = 10^-3^ per generation (for both host and bacteria).

### Stress-dependent adjustment of bacterial niche size

We examined the effects of exposure to a single pulse of toxin, ***T*_*0*_**, applied at ***t*_*0*_** (i.e. **ϑ***(t*_*0*_*)=****T*_*0*_**). On timescales smaller than one host generation (100***Δt***), the bacterial community undergoes selection for less sensitive bacteria, accompanied by a drop in the bacterial population size (Figs. 1A,B). In a system with only one level of selection (e.g. free-living bacteria), this would be the only adaptive change. However, when the bacterial population is symbiotically coupled to a host, the survival of each host and bacterium depends also on the amount of detox secreted by the bacteria (Fig. 1C). The secretion is higher for hosts which react to the toxic stress by increasing their carrying capacity for bacteria (i.e. hosts with **δ** > 0; Supplementary Fig. S1A). This leads to stress-dependent selection of hosts which provide a larger bacterial niche ***K*_*B*_** (Fig. 1D), thus increasing the number of resistant bacteria beyond ***K*_*B*_^0^** (Fig. 1A). The benefit from this increase is two-fold: It alleviates the negative impact of losing bacteria (by assisting recovery of the bacterial population; Fig. 1E, Supplementary Fig. S1B) and increases the total amount of secreted detox (Fig. 1F, Supplementary Fig. S1C). However, when **<*N*_*B*_ *>* _*t*_** is larger than ***K*_*B*_ ^0^**, the benefit from higher detox secretion is accompanied by the negative impact of bacterial overload. The combination of the opposing effects of increasing the bacterial niche size adjusts the size of the bacterial population in a stress-dependent manner which acts to maximize the probability of survival of the host.

**Figure 1:**
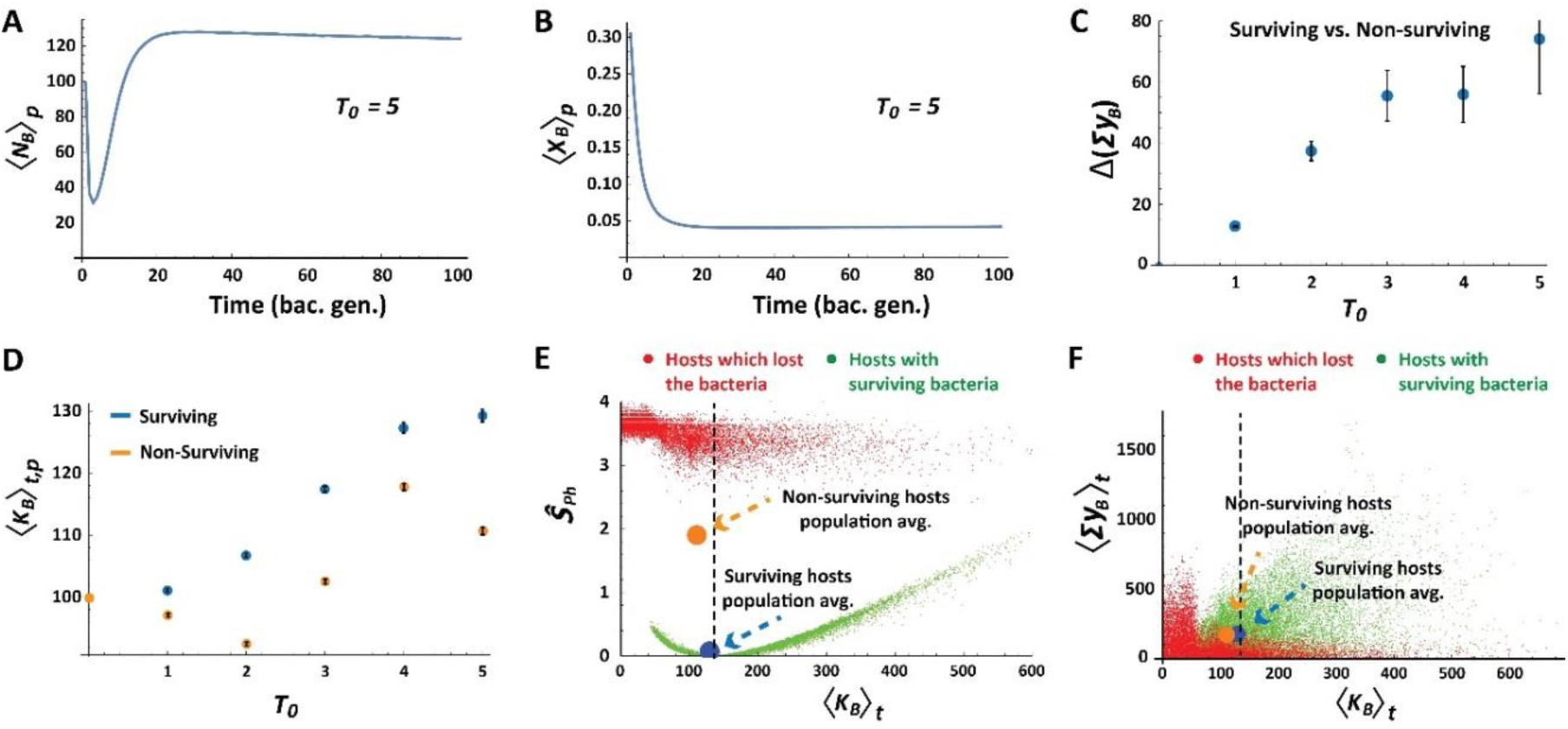
Stress-dependent adjustment of the average bacterial niche size. **(A,B)** Short term kinetics of the population-averaged number of bacteria, <***N*_*B*_**>***P*** (A) and bacterial sensitivity, <***x*_*B*_**>***P*** (B) for hosts which survived a single pulse of exposure to toxin, ***T*_*0*_** =5, applied at the initial time step. **(C)** Average difference ***±*** standard error (SE) between surviving and non-surviving hosts with respect to the total amount of detox secreted by bacteria over a host generation (shown for each ***T*_*0*_**). **(D)** Mean carrying capacity for bacteria in the population of hosts, averaged (± SE) over a host generation at different levels of ***T*_*0*_**. **(E)** Average physiological stress over a host generation, ***Ŝ*_*Ph*_**, versus the time average of its carrying capacity for bacteria. Green and red points represent hosts with surviving and non-surviving bacteria, respectively. Blue and orange circles mark population averages for surviving and non-surviving hosts, respectively. Dotted line marks carrying capacity which minimizes the physiological stress. **(F)** Same as (E) for the time average of total bacterial detox versus bacterial carrying capacity. Time and population averages are denoted by ***t*** and ***p*** subscripts, respectively).

### Stress-dependent adaptation within a host generation

Microbiomes that are modified by the stress in one host generation can be transmitted to the host’s offspring and potentially increase its stress tolerance. In order to evaluate the possibility and magnitude of this outcome we introduce a new measure, termed the “*Lamarckian*”. It quantifies the increase in the survival probability of the host offspring due to (stress-dependent) changes in the microbiome, which occurred during the lifetime of the parental host. To take into account only those changes that were induced by the environmental stress, we compared the survival of offspring hosts to the survival of their parents, judged by exposure at their initial state. To implement this analysis in the simulation, we identify the hosts which survived a generation of exposure, revert them to the initial state of their microbiome and apply a new simulation to the reverted hosts (denoted “cloned parents”) and their offspring (Fig. 2A). We then compare the survival rates of the offspring (*SR*_*Offs*_) to that of their cloned parents (*SR*_*CP*_) and define the *Lamarckian*, ***L***, as:

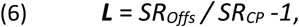

so that it is positive if the average survival increases due to transfer of changes acquired during a host generation. The use of the initial state of the parental host and its microbiome allows us to distinguish the increase of tolerance due to selection of initially better fit parents, from the gain of tolerance due to transmission of changes acquired during a host generation (not present in the initial parental clones).

**Figure 2:**
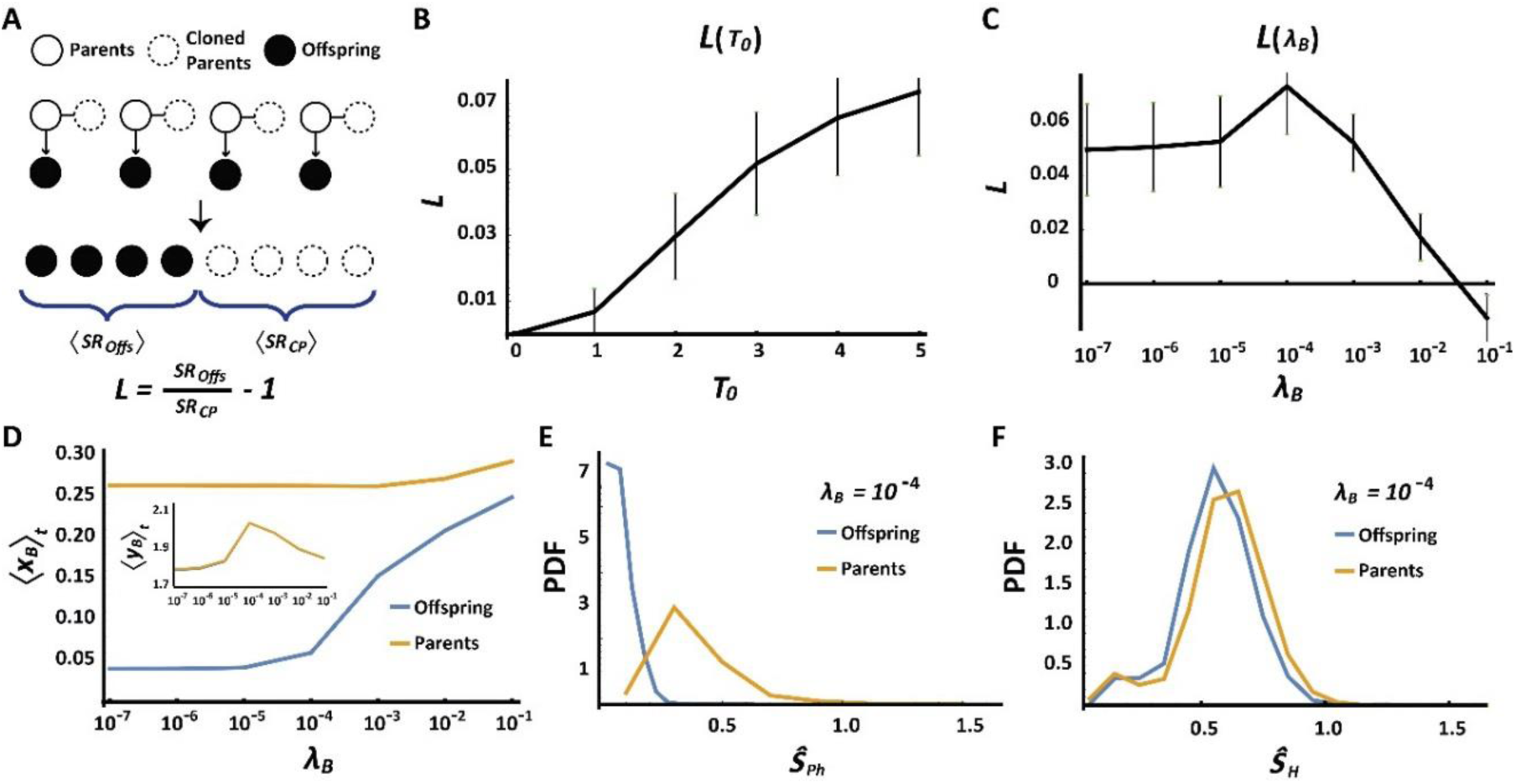
Stress-dependent adaptation within one host generation. **(A)** Schematics of the Lamarckian evaluation protocol. **(B, C)** The *Lamarckian* as a function of toxic exposure (B) and bacterial detox coefficient (C). **(D)** Bacterial sensitivity and detox per bacteria (inset) as a function of bacterial detox coefficient, after exposure to toxin (***T*_*0*_** =5). Shown are time (and population) averages over one generation of unexposed ‘clones’ of surviving parents (orange) and their offspring (blue). **(E,F)** Distributions of physiological (***Ŝ*_*Ph*_**) and toxic stress (***Ŝ*_*H*_**) experienced by cloned parents and their offspring, following exposure to a toxin pulse (***T*_*0*_** =5) applied at the initial time step. Shown for the case of **λ_*B*_** = 10^-4^.

For a given **λ_*B*_**, we find that ***L*** is an increasing function of the injected amount of toxin, vanishing only at low ***T*_*0*_** (Fig. 2B). For a given ***T*_*0*_**, the *Lamarckian* has a non-monotonic dependence on **λ_*B*_**. This is manifested by an essentially constant ***L*** > 0 over a range of small **λ_*B*_**, followed by an increase to a maximum at intermediate values of **λ_*B*_** and lastly, a decline at sufficiently large **λ_*B*_** (Fig. 2C). The positive *Lamarckian* is the result of transgenerational transfer of bacterial population that has been selected for lower toxin sensitivity during the parental host generation (Fig. 2D). To determine how these bacteria increase the probability of survival of the hosts’ offspring, we analyzed the toxic and physiologic stress in the offspring vs. their cloned parents. For small enough **λ_*B*_**, the benefit from bacterial secretion of detox is negligible and the positive *Lamarckian* is primarily due to alleviation of the physiological stress in the offspring (Supplementary Fig. S2A). This is due to inheritance of bacteria that are less sensitive to toxin (Fig. 2D), so that the population size of bacteria in the exposed offspring remains closer to the preferred value (***K*_*B*_^0^**) compared to the bacterial population in their cloned parents. At intermediate values of **λ_*B*_**, the offspring have an additional benefit due to the detox secreted by their toxin-resistant bacteria, thus making a second contribution to the *Lamarckian* (Fig. 2E,F). However, when **λ_*B*_** is large enough to support substantial neutralization of toxin during a single host generation (Supplementary Fig. S3), the selection pressure on both hosts and their microbiomes is weakened and the *Lamarckian* decreases because of the diminished difference between parents and offspring (Fig S2B).

### Selection of hosts based on traits of their bacterial community (‘Microbiome Selection’)

When the toxic pressure persists over timescales larger than one host generation (Fig. 3A), the selection favors hosts with bacterial communities that secrete higher amounts of detox per bacterium, <***y*_*B*_**>**_*P*_** (Fig. 3B). Since this selection is determined primarily by the microbiome as a whole and not by individual bacteria, we will refer to it as *Microbiome selection*. When the secretion of detox comes at a cost to the individual, the microbiome selection for detox is weakened, but it is still apparent over a broad range of cost levels (Supplementary Fig. S4A,B). The negative effect of the cost on the survival probability of bacteria (Supplementary Information, Eq. 2’) aggravates the initial loss of bacteria and increases the physiological stress to the host (Supplementary Fig. S4C). This promotes selection of hosts that can partially alleviate this stress by accommodating larger numbers of bacteria (Supplementary Fig. S4D). The cost on bacterial detox therefore strengthens the selection of hosts which accommodate more bacteria at the expense of weakening the selection for increased detox per bacterium. The Lamarckian effect, on the other hand, is not compromised by the cost of detox (Supplementary Fig. S5A,B), because the increase of physiological stress in parent hosts is larger than the corresponding increase in their offspring (Supplementary Fig. S5C).

**Figure 3:**
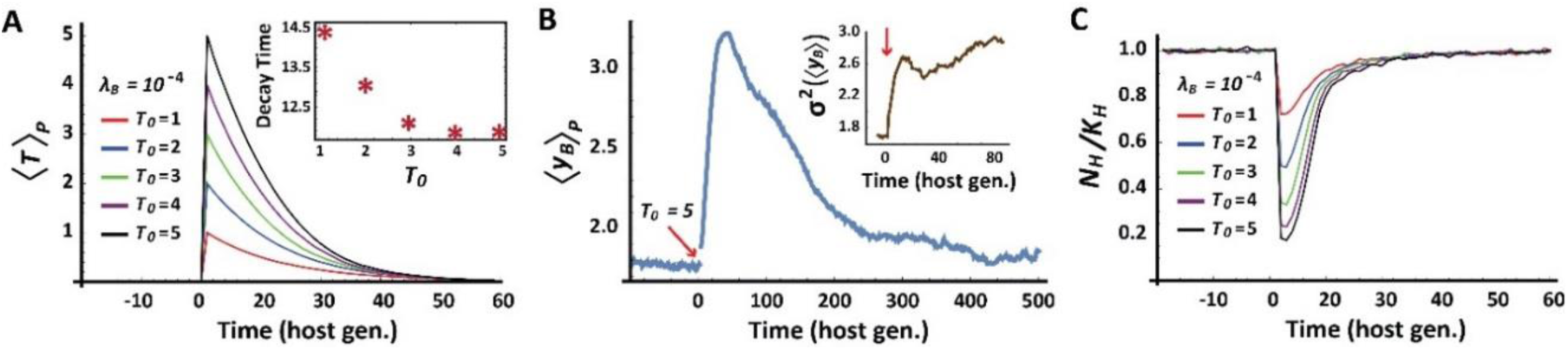
Stress-dependent selection of hosts based on microbiome properties. **(A)** Temporal kinetics of active toxin for different initial levels of toxin, ***T*_*0*_**. Inset displays the time to neutralize 50% of the toxin. **(B)** Temporal kinetics of average detox secretion per bacteria following exposure to toxin at ***T*_*0*_** =5 (red arrow). ***λ_B_*** =10^-4^. Inset reveals an increase of inter-hosts variance in average detox per bacteria. **(C)** Kinetics of host population size, ***N*_*H*_**, normalized by the host carrying capacity, ***K*_*H*_**.

In the current model, the microbiome selection occurs only at the time of host reproduction. If the toxin persists over a period longer than ***µ-^1^*** bacterial generations and the elimination of mutations is sufficiently slow (i.e. small ***β* _*y*_**), the selection is accompanied by significant accumulation of bacterial mutations. Such accumulation enhances the selection for higher <***y*_*B*_**>, thus increasing the detoxification rate (Fig. 3A, inset) and expediting host adaptation (Fig. 3C). This is accompanied by extended persistence of high detox levels (Fig. 3B) and by elevated detox variability across host-microbiome systems (Fig. 3B, inset). Additional increase of variability under stress is observed in the carrying capacity for bacteria and in the size of the bacterial population (Figs. S6A,B).

Following the neutralization of toxin, the selected bacterial mutations persist over a characteristic timescale of 1/***µ*** = 10 host generations, thus providing a ‘memory’ of the previous exposure. To evaluate the influence of this ‘memory’ on the tolerance to new exposures, we analyzed the response to repeated pulses of injected toxin, separated by time intervals shorter than 10 host generations. The re-exposures led to microbiome selections occurring at a rate that is sufficient to oppose the relaxation of <***y*_*B*_**>**_*P*_** to its (lower) equilibrium value (Fig. 4A vs. Fig. 3B). The resulting enhancement of detoxification (Fig. 4B), reduced the selection pressure on the host (Fig. 4C) and enabled the survival of intrinsically less resistant hosts and bacteria (Fig. 4D). Progressive reduction in the intrinsic resistance of the host due to successive selections of higher bacterial detox, is reminiscent of *Genetic Assimilation* by successive selections of host-intrinsic alleles [51, 52]. In the case of microbiome selection, however, the gradual change in the population of hosts is caused by successive selection of variations in the bacterial population (“Bacterial/*Microbiome Assimilation”*). Bacterial variations emerge on faster timescales compared with germline mutations in the host genome, but they are considerably less stable than host-intrinsic mutations. However, when the repertoire of host-intrinsic alleles available for selection is limited, the hosts’ population may become more strongly dependent on variations that emerge within the host’s lifetime (e.g. bacterial and epigenetic variations).

**Figure 4:**
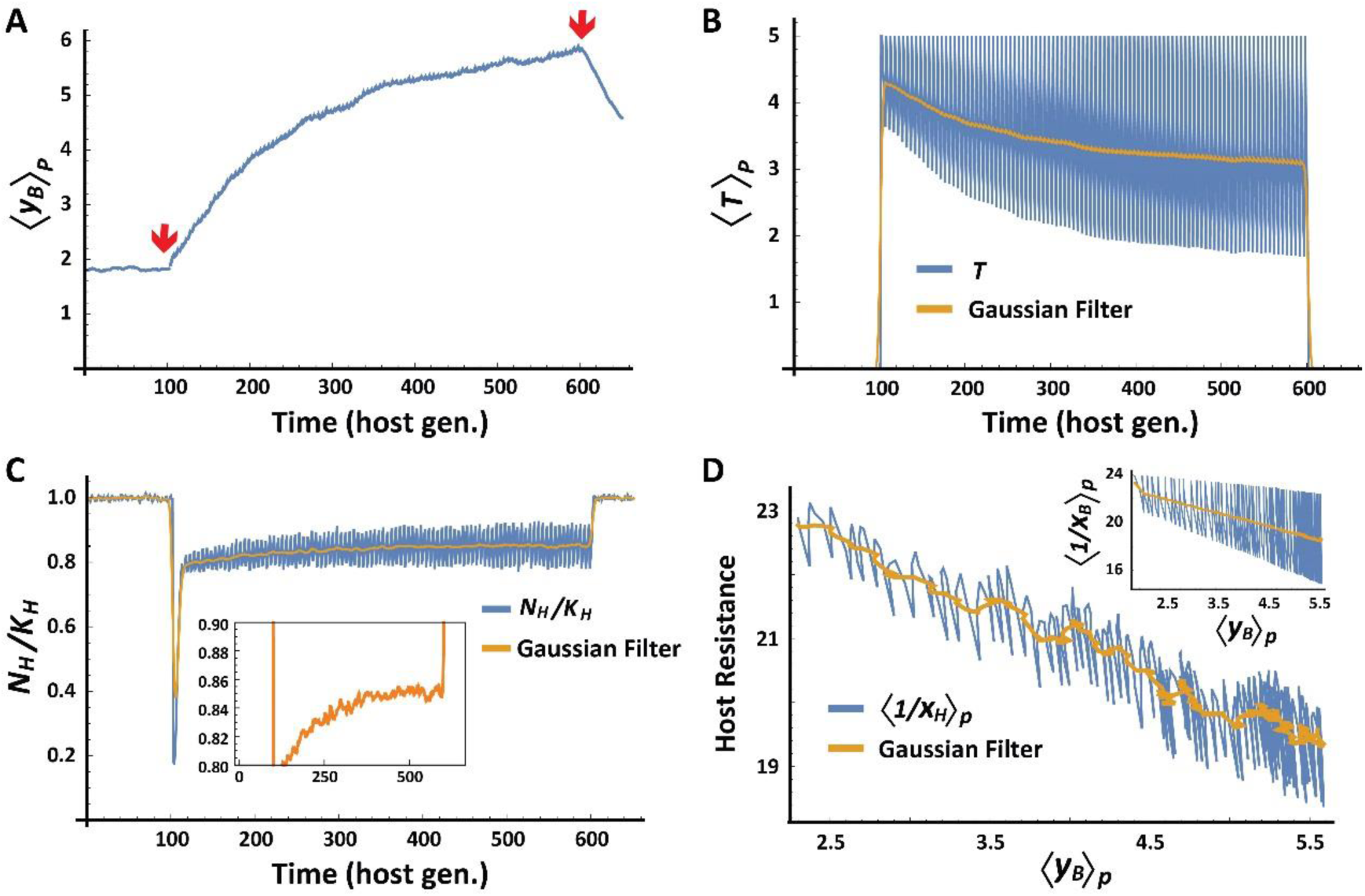
Multi-generational coupling between microbiome properties and host-intrinsic traits. The population of host-microbiome systems was subjected to successive resetting of the active toxin to ***T*** = 5, every 5 host generations. **(A-C)** Temporal kinetic profiles of average detox per bacteria (A), active toxin (B) and normalized size of the host population (C), with a magnified scale in the inset. Red arrows in (A) mark the start and end of the successive resetting of the toxin. **(D)** Inverse correlation between the increase in detox secretion per bacterium and the average toxin resistance (inverse sensitivity) of host, **1/*x*_*H*_**, and bacteria, **1/*x*_*B*_** (inset). Orange overlays correspond to Gaussian filtering of the measured properties.

### Potential strategies for Lamarckian estimation in experimental settings

Quantification of the *Lamarckian* in the model was done by reverting a subset of host-microbiome systems to their initial state and re-subjecting them to toxin. Since we cannot apply this procedure to experimental data, the *Lamarckian* of a real system should be approximated by other means, which may be context-dependent. In organisms such as flies and worms, where the bacteria can be removed without a significant impact on survival (e.g. by egg dechorionation followed by placement on a sufficiently rich diet [53-55]), the *Lamarckian* can be approximated in steps that are conceptually similar to the simulation procedure: first, the hosts are exposed to a challenge and their offspring are cleared of bacteria and separated into two subpopulations. One of these subpopulations is re-colonized with (‘naïve’) microbiota from untreated hosts (as in refs. [34, 56]), while the other is colonized by (‘experienced’) microbiota from a group of hosts which survived exposure to a challenge. The *Lamarckian* is then evaluated using the survival rates of hosts with experienced vs. naïve microbiota (i.e. ***L*** ≈ *SR*_*Exp. microb.*_ / *SR*_*Naïve microb*._ - *1*). This evaluation, however, neglects other types of changes that may have been acquired and transmitted to offspring (e.g. small RNAs [10], maternal RNA [57], persistent chromatin modifications [8], horizontal transfer of biochemical signals [58] or other modes of local niche construction [14], etc.). Additional consideration that may affect the evaluation is horizontal transmission of bacteria to bystander hosts and/or to offspring, which did not inherit the acquired change from their own parents. The above effects can be taken into account by removing the bacteria from two untreated subpopulations, re-colonizing them with ‘naïve’ and ‘experienced’ microbiota, respectively, and estimating the *Lamarckian* from the survival of these colonized populations under challenge. More generally, it should also be possible to obtain a relative measure of the *Lamarckian* by manipulating the microbiome (or any other factor) in a subpopulation of hosts and evaluating the relative difference in offspring adaptation compared to offspring of non-manipulated parents (taken from the same distribution of hosts).

## Discussion

We explored the adaptation dynamics in a host-microbiome model in which Darwinian selection of the host is coupled to a faster selection of its vertically-transmitted bacteria. It is generally accepted that selection of bacteria occurs in every animal and plant and that some of these bacteria can be horizontally and/or vertically transmitted [25, 59, 60]. Transmission of a bacterial population that has acquired changes during a single host lifetime can potentially alter the state of the host and may confer adaptive capabilities that are traditionally considered impossible for germ-free hosts and free-living bacteria. Rigorous evaluation of these capabilities has been hampered, however, by disagreement about how to conceptualize the adaptation and evolution of a composite system of host and bacteria [25, 36-38]. In particular, it is not clear whether the association of the bacterial community with a host (and its offspring) is tight enough to support their co-adaptation and evolution as a (holobiont) unit. Our model bypasses this difficulty by relying on well-accepted Darwinian selections operating, respectively, on hosts and (vertically transmitted) bacteria. We show that interaction between these selections can give rise to previously unrealized modes of emergent of adaptation, promoted by bacterial influence on the survival probability of the host. This includes a gain in offspring tolerance due to toxic exposure of the parental host (Lamarckian effect) and selection of hosts based on attributes of their bacterial communities (Microbiome selection). This was evidenced, for example, by the progressive increase in the host population size (Fig. 4D) despite a reduction in the intrinsic resistance of the host (Fig. 4C).

Within the simplified model in which the survival of the host is evaluated only at the time of reproduction, Lamarckian adaptation arises due to rapid selection and transmission of resistant bacteria. This transmission opposes the loss of bacteria following toxic exposure and confers two types of benefits to the host’s offspring: a) reduction of physiological stress and b) increase in the total detox secreted by the bacteria. The contribution of each of these effects to Lamarckian-like adaptation depends on the level of toxic exposure and the detoxification capacity. Selection of hosts with higher bacterial detox, on the other hand, occurs on a timescale larger than one host generation and therefore cannot contribute to the *Lamarckian* which measures the offspring’s gain in tolerance due to changes that occurred within a single generation of parent hosts. Microbiome selection is nonetheless the main contributor to the progressive increase in tolerance over multiple host generations. Taken together, the transient Lamarckian adaptation is mediated by selection of resistant bacteria within one host generation while the longer-term adaptation under prolonged toxic pressure is achieved by selection of bacterial communities with higher detox per bacterium.

Although the aforementioned capabilities are linked to common features of host-microbiome systems, the scope and generality of the current model are limited by its simplifying assumptions. Studying the effects of factors that are not included in the present work (e.g. multiple species of symbionts and/or pathogens, epigenetic effects, etc.) requires suitable extensions of the model. A noteworthy aspect that is not covered in our model is the potential effect of horizontal transmission of bacteria. While the latter is generally expected to erode specific associations between host and bacteria [61], theoretical analysis of horizontal transfer under selection has demonstrated the feasibility of interspecific epistasis effects even in the absence of perfect transmission [62]. This possibility is further supported by evidence of high interpersonal variability in the composition of microbiota in different body habitats [63-65] as well as by dependence of the microbiome composition on genetic determinants of the host [65, 66] and host-specific factors [67, 68]. Based on the theoretical prediction and the experimental findings (as well as the insights from our simulations), we expect that the selection for higher bacterial detox will be weakened by horizontal transmission, but will vanish only in the limit of strong “mixing” in which all the hosts in a given generation are populated with indistinguishable bacterial communities. Emergent Lamarckian adaptation, on the other hand, should hold even in the extreme case of complete bacterial mixing, because it is mediated by rapid selection of resistant bacteria followed by transfer to the hosts in the following generation. Horizontal transfer is not expected to compromise these processes, but rather to promote sharing of the benefits with other offspring.

The large timescale separation between the selection of individual resistant bacteria and selection of bacterial communities which secrete more detox, reflects a lack of mechanism (in our model) for changing <***y*_*B*_**> during a single host generation (with the possible exception of rare cases of rapid changes in <***y*_*B*_**> due to amplification of very small numbers of resistant bacteria). This limitation can be removed by allowing the stress of the host to influence the distributions of bacterial phenotypes. While we did not consider this type of influence in our simplified model, it likely applies to every host-microbiome system because of the numerous options for 2-way interactions between the host and its symbionts. An extension of the model which allows the stress of the host to influence the bacterial distribution of detox (e.g. by subjecting ***β* _*y*_** to stress-dependent dynamics similar to that of ***K*_*B*_**) could increase the overall secretion of bacterial detox during the lifetime of the host. This could allow the host to benefit from newly-forming bacterial mutations and may further affect the *Lamarckian*.

Finally, we would like to re-emphasize that the proposed modelling framework does not aim to fit a particular host-microbiome system, but rather to investigate the possible modes of adaptation in a system with interactions between selections of host and vertically transmitted bacteria. We show that such interactions can support non-traditional adaptive modes, including a gain in tolerance of the host’s offspring due to the toxic exposure of its parent and longer-term selection of hosts based on collective detox secretion by their bacterial communities. When the toxin persists, or is frequently re-encountered, the selection of detoxifying microbiomes reduces the toxic pressure on the host and weakens the selection of hosts based on their intrinsic resistance.

## Conclusions

Our findings show that interactions between pure Darwinian selections of host and its bacteria can give rise to emergent adaptive capabilities, including Lamarckian-like adaptation of the host-microbiome system. Since the model considers general factors that are typical of host-microbiome systems, the emergent capabilities are likely relevant to most animals and plants and other types of organizations, which satisfy the general assumptions of this modelling framework. The latter can be readily adjusted to incorporate additional factors, such as having multiple species of symbionts and pathogens (with inter-species competition and/or cooperation), asynchronous reproduction modes, epigenetic effects, ecological influences and transfer of bacteria (and/or toxin) between hosts and more.

## Methods

### Simulation procedures

The simulation starts with a population of hosts, each carrying a population of 100 bacteria. Host and bacterial properties (phenotypes) are initially drawn from defined distributions (steady state of Eq. 5 without toxin) with the parameters ***x*_*0*_** = 0.25, ***β* _*x*_** = 10, ***y*_*0*_** = 0, ***β* _*y*_** = 0.1, ***δ*_*0*_** = 0 and ***β* _*δ*_** = 0.1.

In every time step of the simulation (one bacterial generation), each bacterium reproduces if its survival probability (Eq. 2) is larger than a random number (between 0 and 1) drawn from a uniform distribution. Each of the surviving bacteria (parents) persists at its current state and gives rise to a modified bacterium (offspring), while dead bacteria are discarded. At the end of one host generation (100 time steps), the reproduction of hosts is determined based on the survival probability in Eq. 1. Non-surviving hosts are discarded and each of the surviving hosts gives rise to a parent and offspring host as follows:

‐ The parent retains its current state (***x***, ***y***, **δ**) and the state of its bacterial population.

‐ Following 99 bacterial generations, an offspring host is created with properties defined by Eq. 5. Negative values of the sensitivity and detox are prevented by taking the absolute value of the outcome in Eq.1. Each offspring receives a copy of the bacterial population of its parent. These populations are then iterated forward one bacterial generation, the surviving bacteria reproduce so as to define the initial state of the bacterial populations in the next host generation of the parent and its offspring.

## Acknowledgements

We would like to thank Prof. Naama Brenner (Technion, Israel), Prof. Michael Shapira (UC Berkeley). Dr. Michael Elgart (Weizmann Inst. Israel), Maor Knafo (Weizmann Inst. Israel), Prof. Erez Braun (Technion, Israel) and Prof. David Bensimon (Paris ENS-LPS, France) for helpful discussions and suggestions. YS was supported by the John Templeton Foundation grant No. 40663. YR was supported by the Israel Science Foundation grant 1902/12 and the I-CORE Program of the Planning and Budgeting Committee of the Israel Council for Higher Education. DK was supported by the Binational Science Foundation, grant 2015619.

